# Sequoia: An interactive visual analytics platform for interpretation and feature extraction from nanopore sequencing datasets

**DOI:** 10.1101/801811

**Authors:** Ratanond Koonchanok, Swapna Vidhur Daulatabad, Quoseena Mir, Khairi Reda, Sarath Chandra Janga

## Abstract

Sequoia is a visualization tool that allows biologists to explore characteristics of signals generated by the Oxford Nanopore Technologies (ONT) in detail. From Fast5 files generated by ONT, the tool displays relative similarities between signals using the dynamic time warping and the t-SNE algorithms. Raw signals can be visualized through mouse actions while particular signals of interest can also be exported as a CSV file for further analysis. Sequoia consists of two major components: the command-line back-end that performs necessary computations using Python and the front-end that displays the visualization through a web interface. Two datasets are used to conduct a case study in order to illustrate the usability of the tool.

## 1. Introduction

For the longest time, second generation sequencing has been our primary approach to sequence RNA [1]. However, as we dissect the complex nature of RNA more and more, we are learning how short read sequencing does not capture the compound nature of the RNA [2]. Advancements in single-molecule long read sequencing from Oxford Nanopore Technologies (ONT) and Pacific Biosciences [3] are showing promise in developing insights about the RNA role and function.

Especially ONT is expanding its user base rapidly, given its low cost. The MinION from the ONT records the change in the current across the nanopore and uses it to thereby determine the sequence. This consensus of current values is written out as binary hdf5 (DOI: 10.11578/dc.20180330.1) files, accessed under the .fast5 extension files. These Fast5 files are then base called by tools like Guppy to obtain the fastq sequence. The current is sensitive and specific to the moiety passing through the pore. Although there are multiple packages and frameworks built for analyzing and visualizing nanopore sequencing data such as poRe [4], poretools [5], NanoPack [6], NanoPipe [7], and NanoR [8], all of them focus on the whole dataset and do not provide convenient options to look at specific subsets of signals. Most of the existing tools plot statistics about the data rather than providing insights into the data. If the signal can be accurately represented, accessed, and analyzed at much deeper level like for specific k-mers and specific locations; recognizing the functional properties of RNA like mutations, structure variants can be exercised more extensively and robustly.

Especially RNA modifications have recently emerged as critical post transcriptional regulators of gene expression programs. They affect diverse eukaryotic biological processes, and the correct deposition of many of these modifications is required for normal development [9]. With the use of machine learning and deep learning to predict RNA modifications, a visualization tool that can assist in understanding electric features that are broadly associated with modifications will help in developing a new prediction approach. Despite being the need of the hour, tools and frameworks that render such options at the signal level for a given k-mer or a location are next to none.

Therefore, here we present Sequoia, which is a user-friendly visualization tool that allows users to explore signals generated by the ONT in multiple perspectives. It accepts Fast5 files generated by the ONT and then, using the dynamic time warping similarity measure, displays the relative similarities between signals using the t-SNE algorithm. Raw signals can also be visualized through mouse actions, including clicking, brushing, and hovering. Particular signals of interest can be exported as a CSV file for further analysis as well. Sequoia consists of two major components. One is the command-line back-end that performs necessary computations using Python. The other one is the front-end that takes in the output from the back-end as input and displays the visualization through a web interface using a JavaScript library D3.js.

## 2. Methods

### 2.1. Back-end

To process the raw Fast5 output from ONT, we developed HDF5 organization aware adhoc python scripts which extract and reformat the data for further analysis and visualization. All of these steps are automated and sequentially executed in the background when Sequoia is deployed on a dataset. The order in which each of the module in the background is executed is designed to enable memory efficacy while still retaining all the information from the input data. The input file is submitted through a command-line and the output from the preprocessing is written to a local folder which will then be utilized by the front-end visualization. The sequential pipeline of Sequoia is as seen in Figure 1, each block in the pipeline represents the layers of functionality involved in Sequoia, taking raw input data to visualizing it, making it a robust automated end-to-end framework.

**Figure 1.**
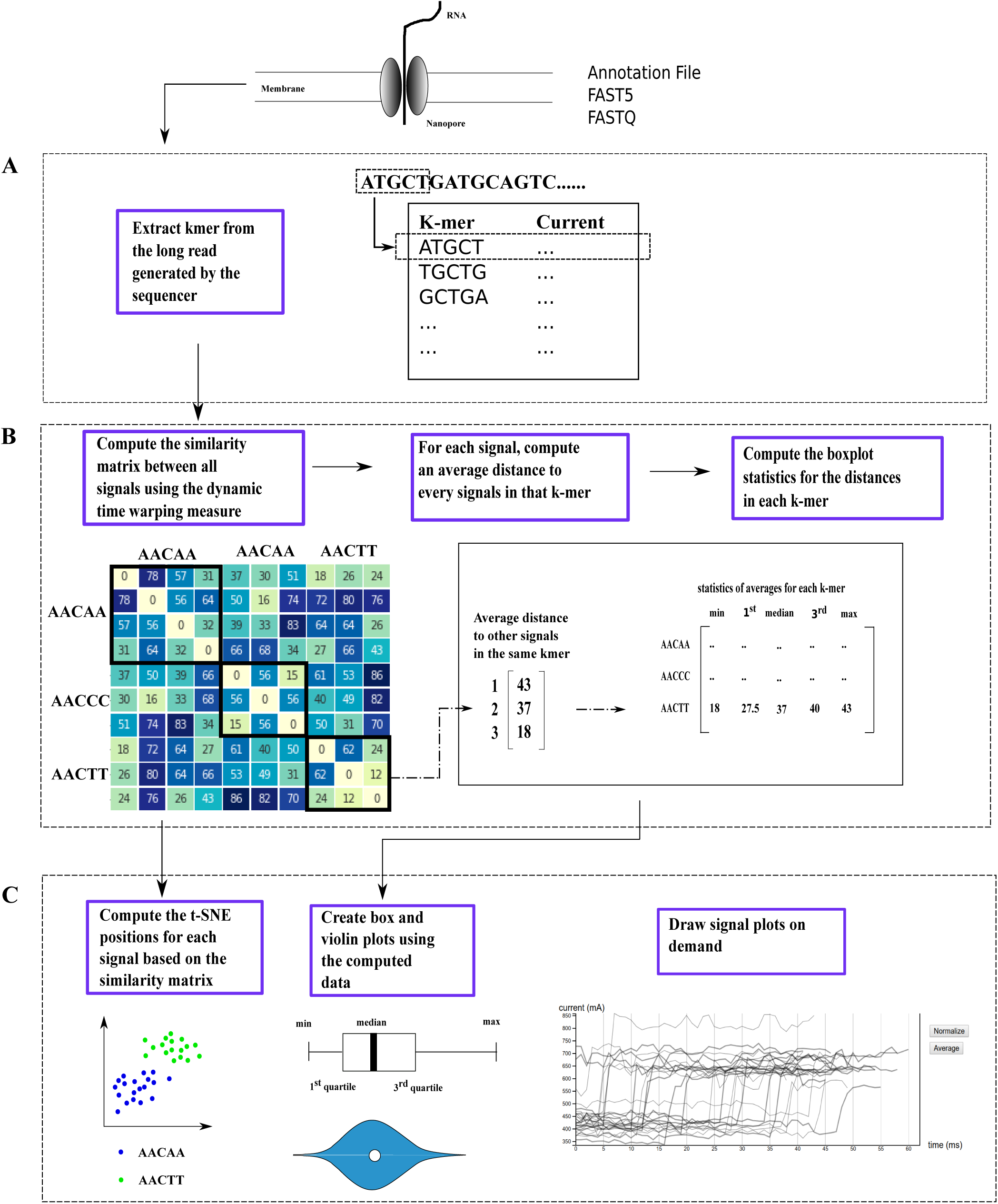
Overall flow of Sequoia (A) Converting long read to a list of 5-mer and their corresponding signal values (B) Computations of dynamic time warping and summary statistics (C) Generating web interface using results from the computations

#### 2.1.1 Input Submission

To parse and extract the signal of interest, the input Fast5 file is read into the pipeline by executing the respective signal extraction script through the command-line. Sequoia renders users to supply four arguments while submitting an input through the command-line: the input file, the dynamic time warping penalty, the 5-mer of interest, and the output directory. The signal extraction script parses through the events table and extracts the respective signal from the Fast5 file, while annotating it to the appropriate 5-mer. The signal annotation process yields a table of consecutive 5-mers and their corresponding signal. After the Fast5 file is processed and converted into a 5-mer signal table, we then perform a series of computations which further processes this signal table and reformats the data in such a way that it is efficiently fed into the visualization pipeline, eventually generating an output folder consisting of similarity matrix, summary statistics, and 5-mer specific tables.

#### 2.1.2 Similarity matrix and dynamic time warping

One of the important features of the Sequoia is that it can visualize relative similarities between signals by utilizing the t-SNE algorithm. The t-SNE algorithm takes in the similarity matrix generated in the previous step to generate the t-SNE scatter plot visualization. Although there are various measures that can be used when computing a similarity matrix, dynamic time warping is a suitable similarity measure for this application, because of the temporal nature of the signals generated by the ONT.

Dynamic time warping is an algorithm for measuring similarities between two temporal sequences that may vary in time or speed.

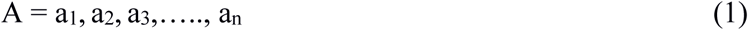

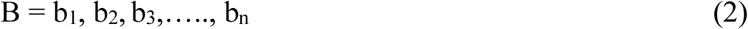

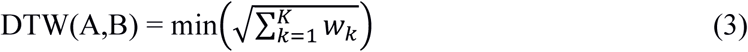

w_k_ is the matrix element belonging to the k^th^ element of the warping path W that represents a mapping between A and B.

The dynamic programming formulation (DTW) as defined above, is based on the following recurrence relation, which defines the cumulative distance for each point [7].

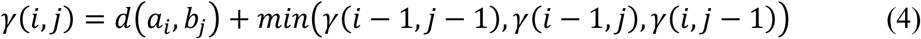

For Sequoia’s back-end, the dynamic time warping computation is performed in python by using the dtaidistance python package (DOI: 10.5281/zenodo.3276100).

#### 2.1.3 Summary Statistics

We used boxplots to visualize the signal distribution and compare it across various 5-mers based on five statistical summaries: minimum, first quartile, median, third quartile, maximum. In order to generate a box plot, these five statistics of the distances for each 5-mer were computed. For each unique 5-mer, an average distance between each signal and all other signals having the same 5-mer sequence was calculated. Then, the minimum, first quartile, median, third quartile, and maximum distances for each 5-mer was computed. The output of this computation is written to a CSV file containing these statistics for each 5-mer.

Similarly, violin plots were generated, which depicts the probability density of the data at different values. For violin plots, the average distances to other signals in the 5-mer group for each signal were calculated. The output of this computation is a CSV file containing average distances for all signals.

#### 2.1.4 K-mer specific tables

In order to visualize the signal plots for specific 5-mers, the raw data is parsed to extract the information only for the 5-mers of interest. After the processing, the single table containing all the 5-mers of interest along with their corresponding signal values will be separated into multiple tables, each containing values from a unique 5-mer only.

### 2.2. Front-end (Visualization)

#### 2.2.1 Tool’s individual functionality

Sequoia provides an intuitive framework for dissecting and visualizing the ONT signal information for all of the 5-mers in the given raw data or specific 5-mers, Figure 2 depicts the various visual demonstrations this tool renders. The followings are details of each individual visualization.

**Figure 2.**
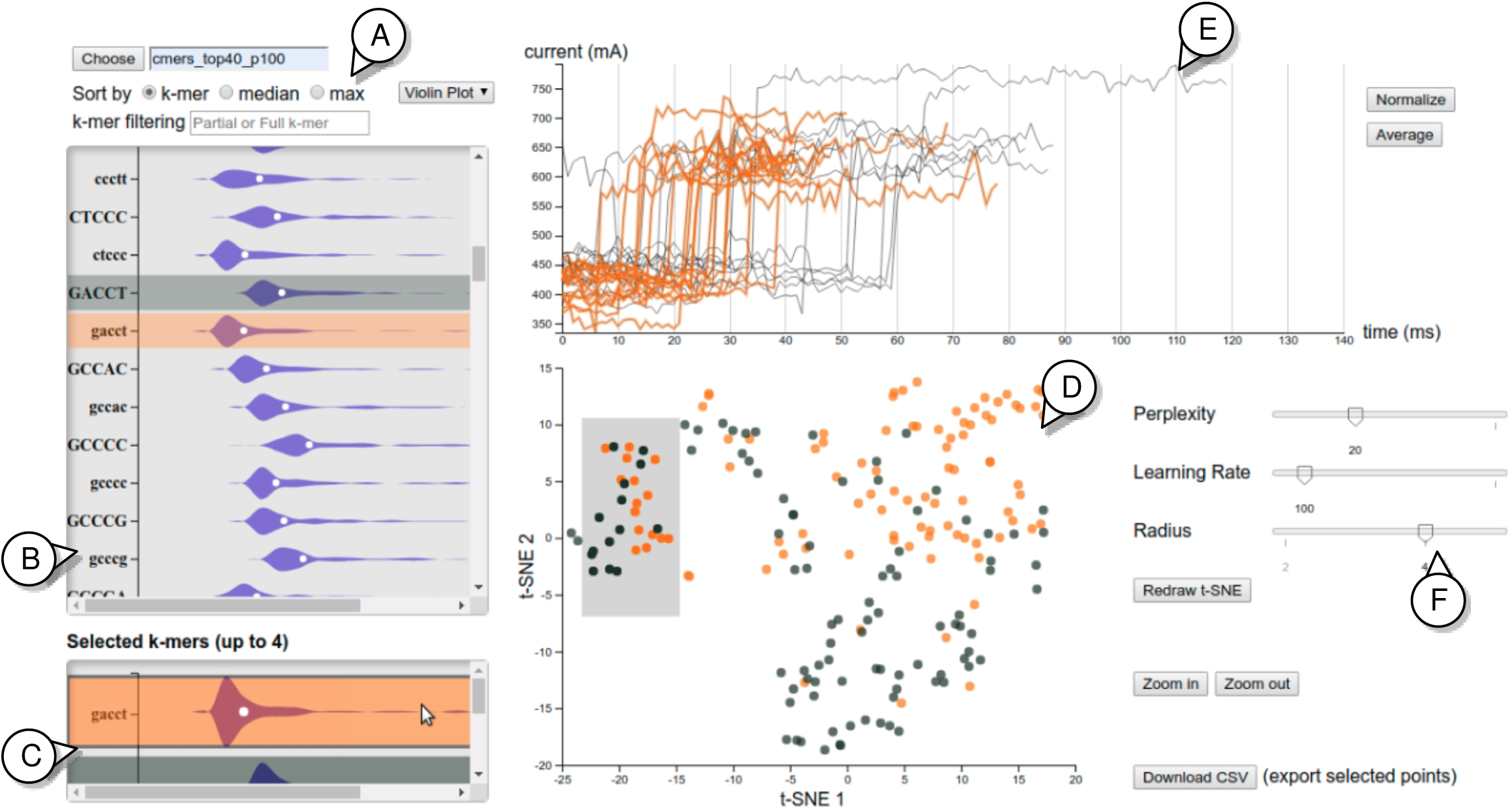
Sequoia’s Interface. (A) Options for displaying 5-mer data including plot types, sorting methods, and text filtering (B) A panel showing a list of all 5-mer in a datasets (C) A panel showing only selected 5-mer from the above panel (D) A t-SNE plot corresponding to the selected 5-mer (E) A signal plot corresponding to the selected points on the t-SNE plot (F) Various options for interacting with the t-SNE plot

##### a) Data Selection (Figure 2A)

By default, the tool is set to visualize data in the directory called ‘data’. If users wanted to visualize data from another directory, they can provide a custom path or folder name into the text box and click ‘choose’. This step makes it possible to submit multiple input data or the same data with different parameter options to the back end simultaneously.

##### b) Box plots and Violin plots (Figure 2A, 2B, and 2C)

Box and violin plots illustrate abstract distributions of the signals corresponding to each 5-mer by displaying the average dynamic time warping distances between each signal and all other signals. The box plots display the minimum, 1^st^ quartile, median, 3^rd^ quartile, and maximum distances between each signal and all other signals on average. The violin plots, on the other hand, provide distributions of the distances. Users can switch between the box and violin plot via a drop-down menu. Users can also sort the order in which the box/violin plot would appear via radio buttons. Specific details are as follows.

- kmer : sort by the k-mer names alphabetically
- median : sort by the median distance among the unique k-mer
- max : sort by the maximum distance among the unique k-mer

Both box and violin plots can be filtered using the ‘k-mer-filtering’ text box. Both plots would exclusively appear based on the user’s sequence of interest typed into the box. For instance, typing ‘A’ would bring up all the 5-mer that start with an ‘A’. In addition, wild card regular expression is also enabled for the text box, a ‘ *’ acts as a wildcard for all four nucleotides (A, C, G, and T). For instance, typing ‘AACA*’ would bring up AACAA, AACAC, AACAG, and AACAT.

##### c) t-SNE (Figure 2D and 2F)

t-SNE algorithm is a technique for visualizing high-dimensional data by mapping data points onto a two-dimensional map [8]. We are using this technique because of its ability to display how close the signals are among the selected 5-mer. Positions for each signal appeared on the t-SNE plot are calculated based on the similarity matrix with the dynamic time warping measure generated in the back end. Once a specific 5-mer is selected, the corresponding t-SNE positions are calculated and the points will be displayed thereafter. Raw data of specific points on the t-SNE can be exported to a CSV file through a ‘Download CSV’ button. The data being exported are read ID, 5-mer, and current value for each signal. Various parameters including the t-SNE’s perplexity and learning rate, as well as the circle size to be displayed, can also be adjusted using the slide bars next to the plot. An option to zoom into a particular region is also available as well.

##### d) Signal plot (Figure 2E)

Signal plots can be displayed on demand in order to explore the similarities between two or more signals explicitly. This view allows users to explore the original signals and compare them side by side. There are two on-demand options that users can select: normalizing and averaging. By default, each signal will be displayed with its original length. A normalization option is available to users for making the length constant for every signal. If the ‘normalize’ button is clicked, the length of all signals will be re-scaled, and thus appear to have identical lengths, as opposed to each having its original length. Similarly, if the ‘average’ button is clicked, the plot will display the average of signals for each 5-mer, as opposed to displaying the original signals.

#### 2.2.2 Interactions across views

Besides having a set of interactions within each view, interactions across multiple views are also possible. Starting from the box/violin plot, up to four 5-mer can be selected through clicking in order to display the t-SNE plot corresponding to the selected 5-mer. Once a 5-mer in the box/violin plot is chosen, two main actions will be performed. First, the box/violin plot of that 5-mer will be highlighted with a distinct color. Then it will appear in the ‘selected kmers’ window. Second, a set of points corresponding to all signals in the 5-mer will appear on the t-SNE area. The colors of the points will be consistent with the colors appeared in the box/violin plot. In addition, mouse hovering over a 5-mer in the ‘selected kmers’ window will highlight the signal plot with the same color.

The t-SNE plot also has the following interactive features that will affect the appearance of signal plots.

- Hovering: When hovering the mouse over a point, the point will turn red, and a signal plot corresponding to that particular point will be displayed also in red. This signal plot will be removed once the mouse is no longer at that point.
- Clicking: When clicking at a point, the signal plot corresponding to that particular point will be displayed in black. This signal plot will be removed if the is clicked again.
- Brushing: Dragging a mouse over a group of points allow a group selection. The squiggle plots corresponding to all the brushed points will be displayed. The plots will change simultaneously corresponding to the changes in brushing action.

### 2.4 Case Studies

To demonstrate the application of Sequoia, we deployed it over two case studies. Each of the case study illustrated the differential nature of the signal from ONT across m5C and m6A RNA modifications. The fore mentioned samples were explored further using Sequoia to understand the nature of the signal from ONT based long read sequencing. Corresponding visualizations were generated to establish the application, scalability and scope of this tool.

#### 2.4.1 Data Generation

##### Cell culture

Hela cell line was purchased from ATCC cell line collection and cultured in DMEM media supplemented with 10% FBS and 0.5% penicillin/streptomycin.

##### Library preparation and RNA sequencing

Libraries were prepared following Nanopore Direct RNA sequencing kit documented protocol (SQK-RNA002). Briefly, total RNA was isolated using Qiagen RNeasy Mini Kit (Cat No. /ID: 74104), followed by PolyA enrichment using Thermo Fisher Dynabeads™ mRNA DIRECT™ Micro Purification Kit (61021). 500 ng of poly (A) RNA was ligated to a poly (T) adaptor using T4 DNA ligase. Following adaptor ligation, the products were purified using Mag-Bind® TotalPure NGS Beads (M1378-00), following NGS bead purification protocol. Sequencing adaptors preloaded with motor protein were then ligated onto the overhang of the previous adaptor using T4 DNA ligase followed by NGS bead purification protocol. The RNA library was eluted from the beads in 21 µl of elution buffer and quantified using a Qubit fluorometer using the manufacturer’s RNA assay. The final RNA libraries were loaded to FLO-MIN106 flow cells and run on MinION. Sequencing runs and base calling was performed using MinKNOW software (Oxford Nanopore Technologies Ltd.) The data output from MinKNOW for Hela cell line was 800k sequence reads using one flow cell. MinKnow generates data as pass and fail folders. FAST5 files from pass folder were again base called using Albacore 2.1.0(Oxford Nanopore Technologies Ltd.) resulting in 500K single molecule reads for Hela cell line which corresponded to full length transcripts ranging from 50b to 8kb.

#### 2.4.2 Data processing

##### Modification location specific signal extraction

To visually compare the ONT generated signal across modified and unmodified RNA bases, we extracted 8737 genomic locations for m5C and 84,149 m6A modifications in HeLa cells from previous RNA modification studies [10–13]. Adhoc scripts previously generated for Raven (https://github.iu.edu/rkadumu/Raven) were repurposed and new functions were appended to complete the location-based signal extraction process. The first step of the signal extraction process was to index the organization of the Fast5 files for the HeLa data generated. This indexing information will be fed into the downstream to efficiently locate the files and navigate the Fast5 more systematically. Simultaneously, we used Guppy to base call the Fast5 data generated and aligned the corresponding Fastq to the hg38 human genome using Graphmap [14]. The resulting sam files were then processed along with the RNA modification genomic coordinates to extract the read index coordinates. All this information was then fed into the in-house developed signal extraction pipeline to extract the signal for a given m5C and m6A modified locations. Similarly, signal for an equal number of random unmodified genomic locations were also extracted for unbiased comparison.

## 3. Results and Discussion

To demonstrate the extent of application of Sequoia we extracted ONT based signal information for two RNA modifications, m6A and m5C to compare them with their respective unmodified nucleotide signals. We also wanted to explore how various parameters dynamically interact and influence the signal visualization.

### 3.1 Case study 1: Signal comparison for m6A vs. unmodified A demonstrates the substantial role of parameters

Sequoia has a diverse set of visualizations to answer specifically to the user defined question. The user can leverage the potential of Sequoia by adapting the appropriate combination of features and parameters. For instance, in this case study we demonstrate how the t-SNE’s perplexity parameter and the signal averaging function aid maximum inference from the raw signal plot.

When using t-SNE to visualize data, the most important parameter to consider is called perplexity, which is the measure of the effective number of neighbors. A perplexity, with a general range of values from 5-50, significantly effects the performance of t-SNE. With low perplexity, the points on the t-SNE plot would be scattered. On the other hand, a larger number of points would be grouped together when perplexity is high. Figure 3A juxtaposes the t-SNE plots of the signals from ACAGG using high and low perplexity. With higher perplexity, a rough signal pattern can be observed, but only two major groups are distinctively separated. When perplexity is adjusted to be lower, the big groups are broken down into smaller subgroups consisting of similar signals. Higher perplexity is more useful when users would like to consider the big picture of the t-SNE plot, while lower perplexity is more effective if users would like to examine local structures, particularly when there are multiple signal shapes within the same 5-mer.

**Figure 3.**
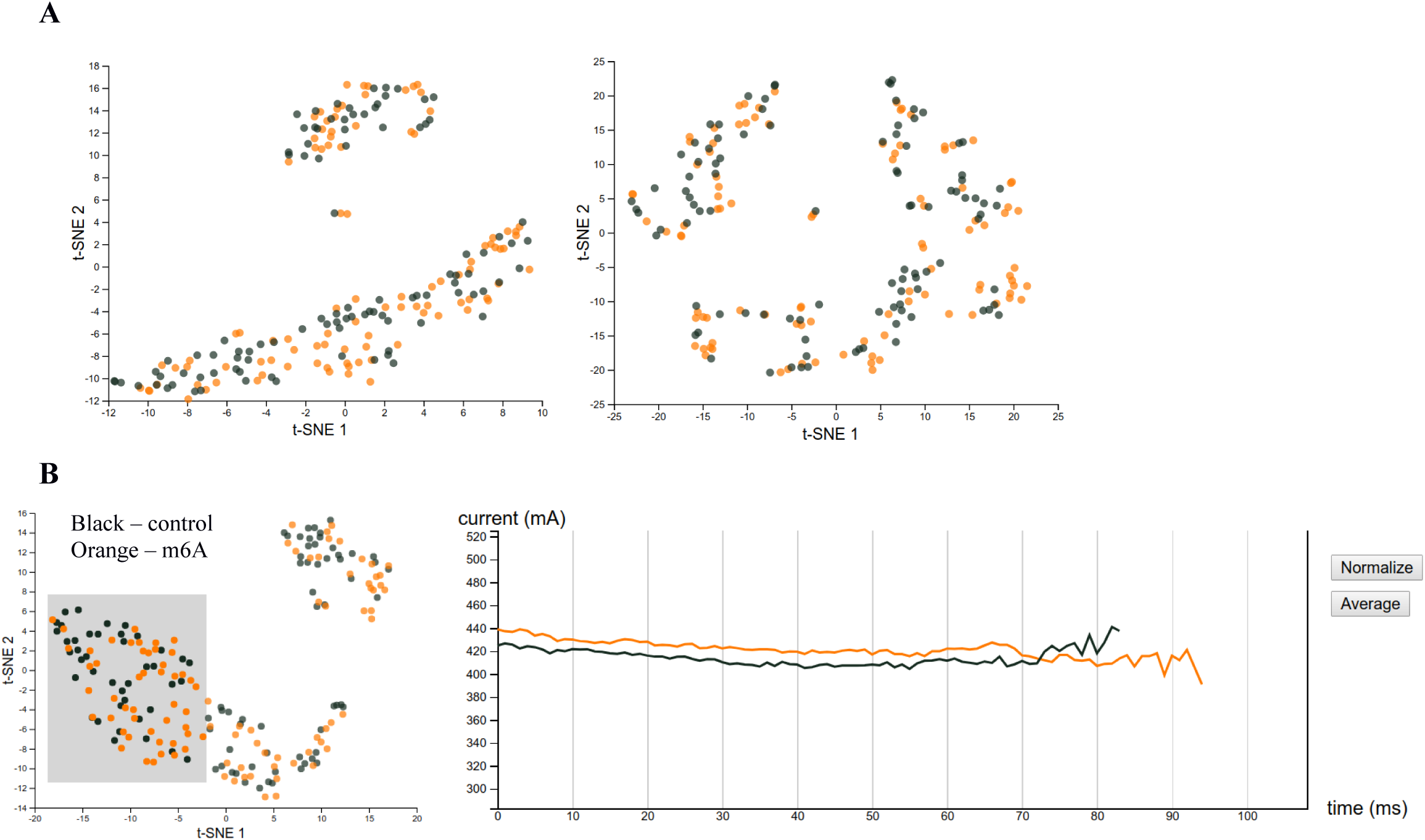
Case study 1 – m6A (A) Comparing t-SNE plots for ACAGG with perplexity = 30 on the left and perplexity = 7 on the right (B) On the right is an average signal plot for ACAGT which corresponds to the selected points in the t-SNE plot on the left. The black line is an average signal of the control group while the orange is an average signal of the m6A group.

Although dynamic time warping and t-SNE parameters can be adjusted, sometimes observing raw signals from another perspective can lead to more conclusive observations. When inspecting raw signals, Sequoia allows users to change the mode from displaying original signals to displaying the average of a selected group instead. This option can help reveal a difference between the unmodified and modified nucleotide signal group that might be overlooked if not for the averages. For instance, figure 3B shows the average signal plot of a group of signals from ACAGT. The t-SNE does indicate that there is some separation between normal and modified signals. The signal plot however shows that the modified group has 20 mA higher current value on average for the selected group relative to the unmodified signal group.

### 3.2 Case study 2: m5C and unmodified C signals illustrate dynamic properties of Sequoia’s visualizations

To demonstrate and emphasize the effect of one of Sequoia’s adjustable parameter called dynamic time warping penalty, which is the distance added if compression or expansion is applied, we compared the signals extracted from m5C and unmodified C in HeLa cell line. The two datasets used in this study were from the m5C data consisting of 19,391 5-mer with modified ‘C’ in the middle and the unmodified data consisting of 537,697 5-mer with unmodified ‘C’ in the middle. In this study, we focused only on unique 5-mers that had at least 100 signals in the m5C dataset. Rest of the data was filtered out, with a final residue of 46 unique 5-mers. Further, for each of the 46 5-mers, we randomly sampled 100 signals from each dataset to be compared in the visualization.

After the preprocessing, we fed the data into Sequoia’s backend and generated the corresponding visualization. In the first attempt, the dynamic time warping penalty was set to zero. After visualizing the data, we decided to focus on GACCT since it was one of the 5-mers that best showed a promising visual separation between normal and modified signals on the t-SNE plot. It can be seen in the left plot of Figure 4A that the points with homogeneous colors are locally grouped together. Although the initial result was satisfactory, it would have been better if we were able to make an improvement so that the separation between groups was more distinct. This led us to start digging into the visualization with the hope of achieving useful insights.

**Figure 4.**
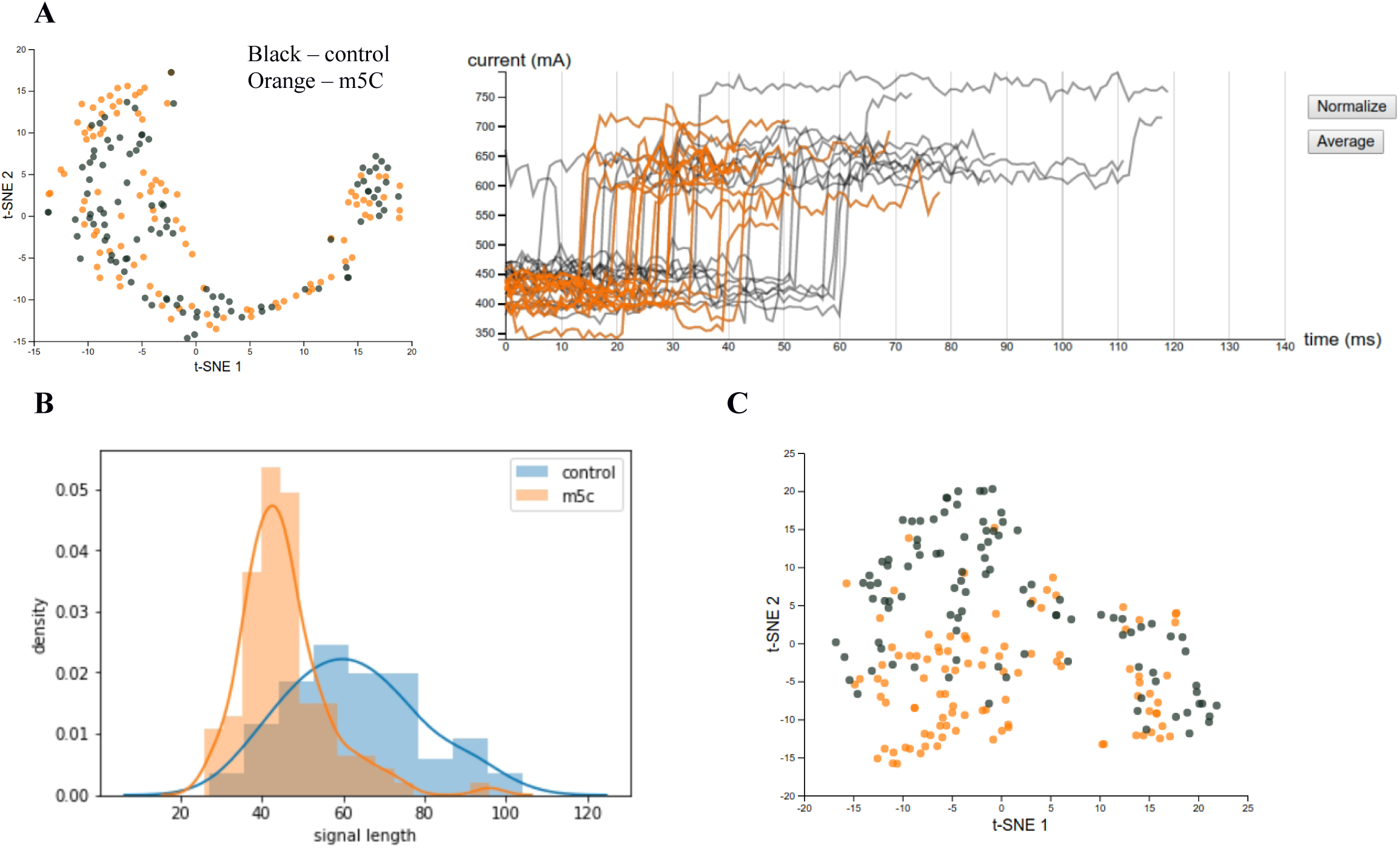
Case study 2 – m5C (A) Left: t-SNE plot for GACCT with no dynamic time warping penalty applied. Right: Raw signal plot corresponding to the highlighted points in the left figure (B) Comparing signal length distributions for the control and m5C signals of GACCT (C) t-SNE plot for GACCT after applying dynamic time warping with penalty

To explore the data further, we utilized the brushing tool for group selection, which is one of the functionalities provided by Sequoia. By brushing over a group of points on the t-SNE plot (as shown in Figure 4A), we were able to see the raw signals plots of those corresponding points along with their corresponding colors. We observed that for the GACCT signals that we had sampled, there was a noticeable difference in signal lengths between the normal and modified groups. To confirm that the length difference noticed was truly significant, we used the Welch’s t-test to test our sampled data with the null hypothesis that the two populations have equal means. The resulting p-value turned out to be less than 0.001, implying that the population means of the two groups were not equal. The distribution plots for both groups are shown in Figure 4B.

After confirming the significant difference in signal lengths of the two groups, we regenerated the visualization by cooperating this new knowledge. Given that the signal lengths were largely related to the compression and expansion of signals during the dynamic time warping process, the dynamic time warping penalty was now set to 100, as opposed to 0 in the previous attempt. As a result, the t-SNE plot for GACCT successfully displayed a much better separation between points representing normal and modified signals, as shown in Figure 4C.

## 4. Conclusion

Sequoia is a tool that can visualize nanopore sequencing data in detail and help biologists discover patterns, develop questions and make inferences. This framework can be used to capture differences between unmodified and modified signals as illustrated in the two case studies. Case study 1 shows how a t-SNE parameter called perplexity and an averaging function in the signal plot can be utilized to exhibit data from different perspectives. Case study 2 demonstrates how an insight obtained from an initial data investigation using the tool can be used to successfully improve the visualization of signal separation between two groups.

